# Emergence timing and voltinism of phantom midges, *Chaoborus* spp., in the UK

**DOI:** 10.1101/676874

**Authors:** R. Cockroft, W.R. Jenkins, A. Irwin, S. Norman, K.C. Brown

## Abstract

After introduction of overwintered fourth instar larvae (2027 in total), emergence timing of adult *Chaoborus* spp. (Diptera: Chaoboridae) was investigated in four outdoor freshwater microcosms in the UK in 2017. Adults started emerging on 13 April and emergence reached a peak on 2 May. The majority of emergence was completed by 3 June. Emergence rates for each microcosm ranged from 51.4% to 66.2% with a mean of 60.9%. The great majority of emerged adults were *C. obscuripes* (99.68%). Males appeared to emerge slightly earlier than females. The results indicated that for overwintered *C. obscuripes* larvae, the adults emerged en masse in spring (rather than emerging gradually over the course of spring and summer). In a separate experiment at the same location, the number of *Chaoborus* spp. life-cycles occurring per year was determined using six replicate groups of microcosms, each group containing four microcosms. Each microcosm contained 200 L of water and was enclosed within a ‘pop-up’ frame covered with ‘insect-proof’ mesh (1 mm^2^ aperture). The first microcosm in each group was ‘seeded’ with egg rafts (first generation) of *Chaoborus* spp. Following adult emergence, as soon as the first egg rafts were laid in each microcosm these were removed and transferred to the second microcosm in that group, and so on. The larvae sampled from the second and subsequent generations in the microcosms were all *C. crystallinus*. *C*. *crystallinus* produced up to four discrete generations within the experimental period, and life-cycle times from egg-to-egg ranged from 14 days (replicate group 5, first generation) to 56 days (replicate 3, second generation). These two experiments, indicated that i) adult *C. obscuripes* arising from overwintered larvae emerged en masse in the spring, and ii) up to four generations of *C. crystallinus* occurred; i.e. *C. crystallinus* exhibited a multi-voltine life history under the temperate conditions of this UK study.

## Introduction

The precise duration of the life-cycle of *Chaoborus* spp. does not appear to have been clearly defined in published studies. Since the larval stages are known to be extremely sensitive to the effects of insecticides (1) and size-dependent sensitivity can play an important role in the survival and recovery of natural populations, the duration and timing of the life cycle has implications for the interpretation of how natural populations recover from exposure to stressors.

The Chaoboridae are Diptera with aquatic larvae that live as pelagic predators, feeding on a wide range of prey including copepods, Cladocera, rotifers, chironomids, mosquito larvae and other chaoborids. There are three genera of Chaoboridae in northern and central Europe; *Chaoborus*, *Mochlonyx* and *Cryophilia*, all of which have four aquatic larval instars (2, 3, 4). *Chaoborus* species are widely distributed throughout Europe and of the six species that are known *C. flavicans*, *C. obscuripes* and *C. crystallinus* are the most common, the most abundant, and the most studied. Pelagic third and fourth instar larvae of some species, such as *C. flavicans*, have adapted to co-exist with fish in larger water bodies and lakes. *C. flavicans* exhibits diel vertical migration to sediment where they burrow during the daytime to avoid predators and migrate upwards at night to feed. In shallower water, or where fish are absent, they may be entirely pelagic (5). Populations of *C. crystallinus* predominantly occur in shallow water bodies without fish and are mostly pelagic, although larvae have been found in sediment (5). Females of *C. crystallinus* are able to detect the presence of fish kairomones in water (6) and so can avoid depositing their eggs in water bodies containing fish.

Larger natural or anthropogenic water bodies that do not contain fish have been found to be dominated by *C. obscuripes* (7, 8). Habitat preferences for the more common European species of Chaoboridae are summarised in Table 1.

**Table 1:** Habitat preferences for the larvae of European Chaoboridae

| Species | Location and Habitat |
| --- | --- |
| *Mochlonyx velutinus* | Larvae found in puddles, springs, pools, wells, tree holes and in temporary or permanent pools. |
| *Chaoborus flavicans* * | Lakes most often but also ponds. |
| *Chaoborus crystallinus* * | Shallow ponds without fish, shallow ephemeral water bodies. |
| *Chaoborus obscuripes* * | Man-made waters bodies without fish, small and shallow waters, also found in deeper lakes. |
| *Chaoborus pallidus* * | Shaded portion of pools. |

\* overwinter as instar IV larvae in permanent water bodies

Chaoboridae are holometabolous and larvae develop through four instars and then pupate. *Mochlonyx*, *Cryophilia* and *Chaoborus nyblaei* overwinter in the egg stage which is resistant to desiccation whilst other temperate species overwinter as fourth instar larvae and pupate between April to June, depending on temperature.

Mating and oviposition take place a few hours after emergence and in some species (e.g. *C. flavicans*) eggs can be held at the surface by a surrounding jelly. *C. crystallinus* lays 200-300 eggs (8) arranged in the form of floating discs to form a raft. The duration of the egg stage is temperature-dependent and for *C. crystallinus* can range from 190-200 hours at 10 ºC and between 37-50 hours at 20 ºC (2). First and second instar larvae of *C. crystallinus* develop rapidly over a few weeks whereas the developmental periods of the third and fourth instar larvae are considerably longer. First and second instar larvae are positively phototactic (9, 2) at first and stay in the upper layer of water (epilimnion), which is warmer and has more oxygen. The later instars are generally found deeper in the water column where they feed on zooplankton.

*Chaoborus* larvae are very tolerant of a wide range of unfavourable environmental conditions. *C. flavicans* can withstand periods of up to 18 days without oxygen when the surface could not be reached and 70 days with access to the surface (2). Larvae of the same species can have variable life spans (10) depending on environmental conditions and the structure of the community. Life cycles can therefore be univoltine (predominantly in Europe) bi-voltine (high temperatures in Europe) or multi-voltine in Japan (11).

The duration of the pupal stage is also temperature dependent and for *Chaoborus crystallinus* this can range from 2-4 days at 20 ºC, between 10-13 days at 10 ºC and 30 days at 5 ºC. In Central Europe, there appears to be a very pronounced emergence period for *Chaoborus* and *Mochlonyx* between April and May with a second, less pronounced emergence from the end of July to October (10). Females emerge from pupae and mate almost immediately, before their genitalia harden. Male swarming behaviour is commonplace and some species have been shown to be attracted to light. Imagines have no resting periods and live for at most ten days during which they do not feed (6, 9).

The presence of larvae throughout the year has led to the conclusion in published literature that *Chaoborus* spp. are univoltine in temperate conditions. In central Europe, *C. crystallinus* was considered to be univoltine, although possibly bi-voltine in hot summers (9, 12–16). Verberk et al (16) considered the dispersal strategy of *Chaoborus crystallinus* to be uni/bi-voltine and typical of those species that have a long period of juvenile development. Life cycle strategies of *Chaoborus* spp. reported in the literature are summarised in Table 2.

**Table 2:** Number of generations per year for *Chaoborus* spp. from published literature

| Species | Number of generations |
| --- | --- |
| *C. flavicans* | Univoltine (8, 18, 19, 20)<br>Univoltine or bivoltine (17) |
| *C. crystallinus* | Univoltine (2, 8, 9, 13, 15, 16, 21)<br>Univoltine and possibly bivoltine in hot summers (12)<br>Multi-voltine (22, 23) |
| *C. obscuripes* | Multi-voltine (24)<br>Univoltine (25) |

Berendonk and Spitze (12) state in their introduction: ‘*Chaoborus crystallinus* is univoltine in Central Europe although it may go through two generations in exceptionally hot summers.’. However, no citation to support this statement is given in that paper. The *Chaoborus* found in microcosm studies conducted in central Europe appear to be mostly *C. crystallinus*, as deduced by Janz et al. (23), who analysed data collected from 19 microcosm studies conducted over 14 years at the University of Munich, Germany. In that study, 1^st^ and 2^nd^ instar larvae were present from mid-April to early October, 3^rd^ instar larvae from early-May to October/November. Larvae overwintered as 4^th^ instar and these were present during the entire study. Pupae were found from early-April to the end of August. The number of egg-laying peaks was used to indicate the number of generations of *C. crystallinus* in that study (the first at the end of April/early May, the second in June and the third at the end of July beginning of August). Janz et al concluded that there were three generations of *C. crystallinus* each year.

As shown in Table 2, publications up to 2008 have considered *C. crystallinus* to be univoltine. On the other hand, the two most recent papers (both published in 2016) are unequivocal in the conclusion that *C. crystallinus* is multi-voltine (in temperate conditions). We would surmise that the disparity between the earlier and most recent articles is that the conclusions of the latter are based on empirical evidence, whereas the statements on voltinism in the other papers appear to originate from an assumption in earlier publications. The empirical evidence in (23) and (23) is essentially the observed prevalence of the various *C. crystallinus* life-stages over time in outdoor microcosms. The experiment described in this publication is a refinement of this approach, directly tracking the progress of successive generations. The difference is that this new work excludes the confounding factor of egg deposition by adults which have emerged from other water bodies. This was achieved by the enclosing microcosms in ‘tents’ made of ‘insect-proof’ netting. The only *C. crystallinus* inoculants were egg rafts placed by the experimenters. These rafts came from the previous (also enclosed) generation. This could be described as a ‘temporal chain’, each link in the chain being an artificial transfer of egg rafts from one enclosed microcosm to the next.

*C. crystallinus* is considered to be highly sensitive to the effects of certain insecticides and has been used in individual based models to predict their potential effects and recovery of aquatic invertebrates (26). An evaluation of the number of generations of *C. crystallinus* per year would be relevant for understanding the recovery potential of *Chaoborus* spp. populations in freshwater systems following possible reduction by pesticides.

## Materials and Methods

Two separate trials were conducted at AgroChemex Environmental Ltd., Aldhams Farm Research Station in Essex, U.K. (Grid Reference TM 099 305) in 2017, one to investigate the emergence timing of *Chaoborus spp.* adults from an overwintered cohort of 4^th^ instar larvae and one to determine how many generations per year could occur in the UK.

Microcosms of the same design were used in both trials. Each microcosm consisted of a circular polypropylene tank approximately 0.8 m in diameter and 0.6 m deep (with a volume of 227 litres), sunk into turfed ground to a depth of approximately 50 cm. 20 litres of washed sharp sand was added and each microcosm was filled to a depth of approximately 50 cm (the same level as surrounding soil) with approximately 200 L of freshly-drawn borehole water. Approximately 10 litres of water-saturated lake sediment were poured in and allowed to settle in an even layer over the sand base and each unit was covered with insect-proof mesh (1 mm^2^ aperture) to prevent the entry of aquatic flies, particularly *Chaoborus* spp. An oxygenating submerged aquatic macrophyte (*Elodea canadensis*, sourced from Envigo, Eye Research Laboratory, Suffolk, UK) was loosely planted in each microcosm to occupy an area of approximately ¼ of the sediment surface. The plant material was rinsed thoroughly prior to introduction into the systems to remove any invertebrates.

Populations of zooplankton (typically comprising rotifers, copepods, daphniids to provide a food source for Chaoborus larvae) and detritus-feeding benthic invertebrates (e.g. *Asellus* and *Gammarus* to facilitate natural recycling of nutrients) were sourced from a natural pond on the site at Aldhams Farm Research Station and added to each microcosm. A handful of alder (*Alnus glutinosa*) leaves was also added to each microcosm to provide a substrate for benthic invertebrates. Alder leaves had originally been collected from Fen Alder Carr (a local nature reserve established in 1982), Suffolk, UK, and then dried and stored. The alder leaves were soaked for >7 days in clean borehole water and roughly shredded with scissors prior to addition to the microcosms.

Active microcosms in both trials were monitored weekly for temperature, pH, dissolved oxygen and conductivity using a Hach HQ40d portable multimeter. Water temperature in one unit was monitored with readings every 30 minutes, using a calibrated data logger. Additionally, the water temperature in an unused, unenclosed microcosm was also monitored continually from June onwards, to allow a comparison of temperatures in enclosed and unenclosed systems. Climatic conditions on the microcosm site were recorded throughout the study using a Davis Vantage Pro2 Plus field weather station, situated approximately 100 metres from the study area.

### Emergence timing

This study was conducted to determine whether the first spring emergence of adult *Chaoborus* from overwintered 4^th^ instar larvae took place over a defined period, or was protracted, with adults emerging at intervals throughout the year.

Four microcosms established between February and March 2017 were used for this study. Populations of *Chaoborus* spp. were established on 22 March 2017 in each microcosm by the addition of approximately 500 4^th^ instar larvae, obtained from an untreated field reservoir at Envigo, Wooley Road, Alconbury, Huntingdon, Cambridgeshire, UK. *Chaoborus* were collected using a sweep net and transferred to a covered holding vessel containing water from the source reservoir for transportation to the field site. Each container held larvae collected from several sweeps of the water column, just below the water surface. On arrival, larvae were held outdoors in their original containers with loosely fitting covers. On the day of initiation, groups of approximately 50 larvae were transferred into a tray, counted and then added to one of the replicate microcosms. This process was repeated until the four microcosms contained 502, 503, 510 and 512 larvae respectively. Before the start of the study, the microcosms were covered with insect-proof netting to prevent colonisation by the local populations of *Chaoborus*. Plate 1 shows the caged microcosms and Plate 2 shows an adult male *Chaoborus* spp.

**Plate 1.**
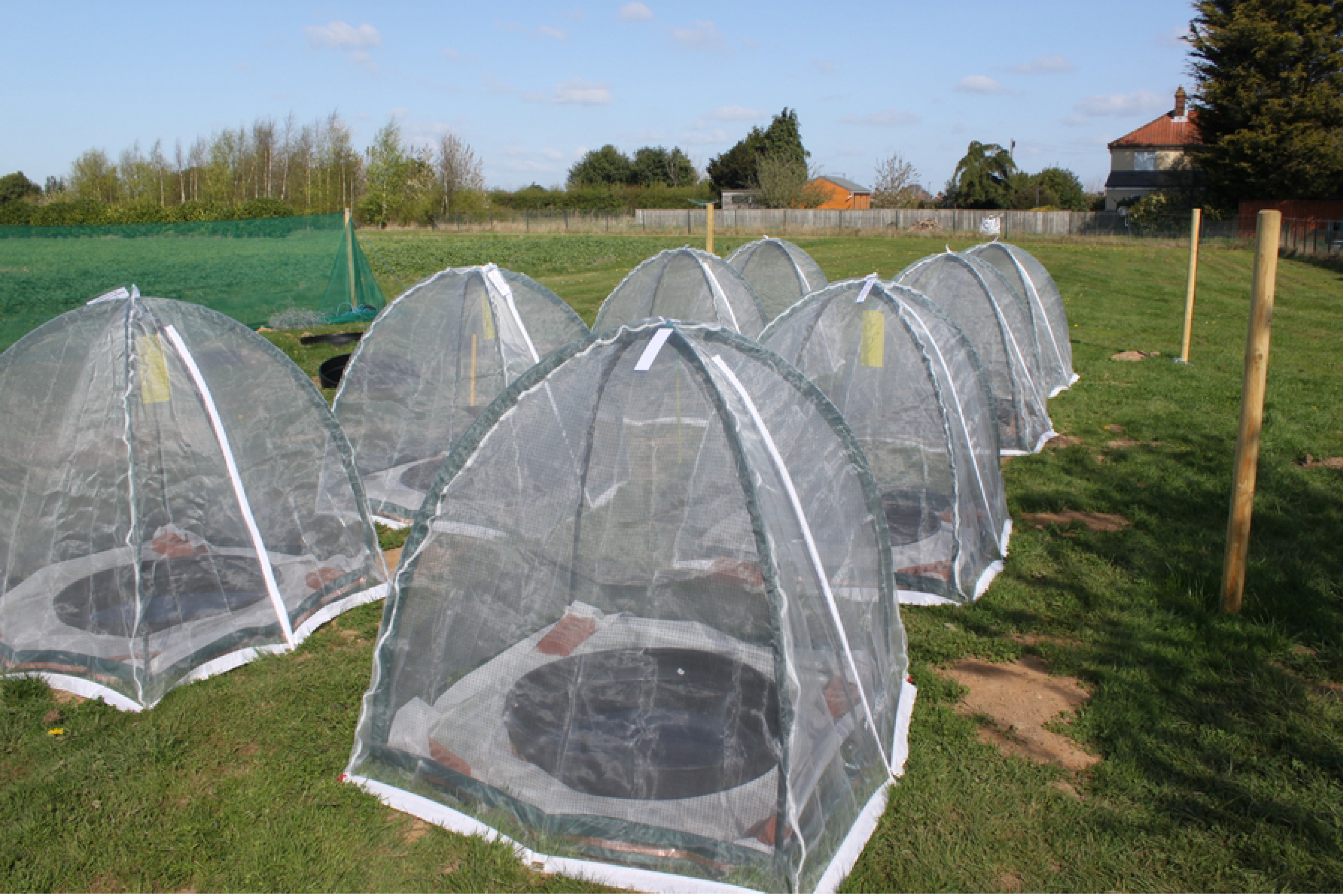
Microcosms with mesh-covered frames.

**Plate 2.**
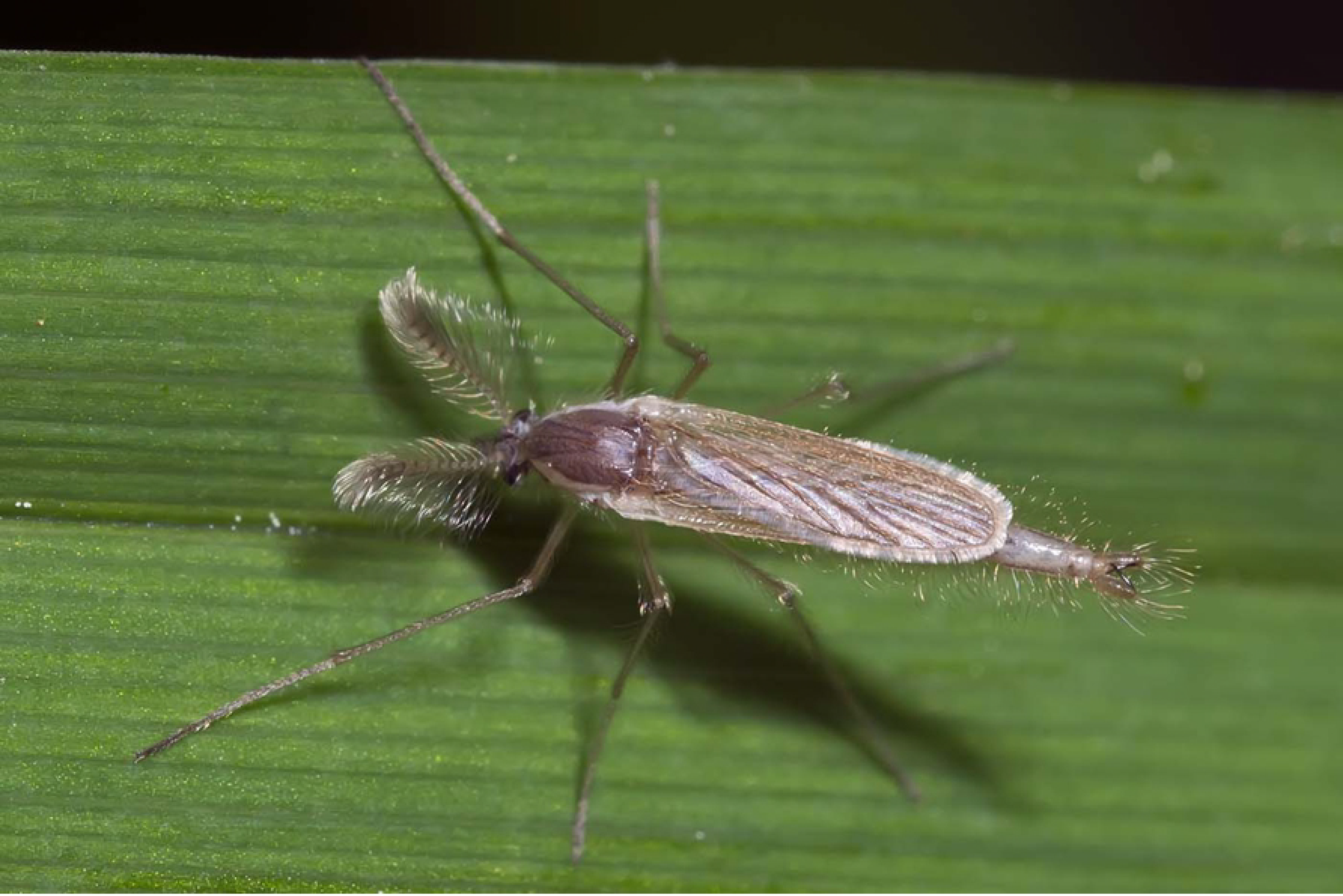
Adult male *Chaoborus crystallinus* male. Photo Eugène Vandebeulque

The emergence of adult *Chaoborus* sp. was monitored daily in the four microcosms by collecting emerged insects seen on the walls of the enclosure or on the surface of the ground around the microcosm using a vacuum sampler. Collected insects were preserved in 70% alcohol and stored for subsequent identification to species level. The water surface was inspected daily for the presence of egg rafts deposited by emerged females that had mated before sampling. Any egg rafts observed were removed as soon as practically possible to prevent the introduction of fresh larvae to the microcosms.

### Voltinism in *Chaoborus*

This experimental system consisted of six replicates each of four individual microcosms established between February and March 2017 at the field site. An additional set of four microcosms was established to provide the initial egg rafts and these were covered with a pop-up frame and insect proof mesh (Popadome, Harrod Horticultural, Lowestoft, UK).

Each of the four microcosms established for the production of egg rafts were initiated with approximately 500 4^th^ instar *Chaoborus* spp. larvae from the same source as used to initiate the emergence experiment and covered with insect-proof netting. Following the emergence of adult *Chaoborus* spp., these egg generation microcosms were regularly monitored for the presence of egg rafts on the water surface. When egg rafts were found they were transferred to the first of the four microcosms in each replicate set. The production of egg rafts in the egg generation microcosms was monitored until no more egg rafts were required. Each of the first of the four microcosms in each replicate set containing egg rafts was inspected at least three times each week for the appearance and development of larvae, pupation, emergence of adults and deposition of egg rafts. The presence of larvae and their approximate instar (estimated by eye) together with the presence or absence of pupae and emerged adults was recorded by inserting a 19 cm diameter white disc attached to a rod to provide contrast for assessing both at the surface (early-instar larvae) and at depth (late instar larvae and pupae) and visual inspection of the enclosure mesh (adults). The date and numbers of any egg rafts produced were also recorded.

Emerged adult insects were allowed to remain within the enclosure, reproduce and deposit egg rafts on the water surface. These eggs were then collected and added to the second microcosm of each replicate set to initiate populations. Established microcosms were inspected three times per week until the end of September when the monitoring frequency was reduced to once per week. The presence or absence of each life stage of *Chaoborus* was recorded in each active microcosm. Where present, an approximation of the size range of larvae visible was recorded, mainly to facilitate monitoring of egg hatching success and the rate of development, to ensure that critical development stages were not missed. Once adult emergence had been observed, at each subsequent assessment, the water surface was inspected for the presence of egg rafts deposited by emerged females. The observation of the first deposition of egg rafts was recorded and those egg rafts used to initiate the next sequential unit within the replicate. Subsequently, additional egg rafts produced within the active units were also transferred to supplement the next unit’s population, until it was considered that no more were required.

On three occasions, once in July and twice in October, samples of late-instar larvae were taken and preserved in 70% alcohol for identification to species level. In the July sampling, only three microcosms contained larvae considered sufficiently developed for identification and ten larvae were sampled from each. In October, where larvae were abundant, approximately 30 were sampled and if fewer than this were seen, all larvae which could be captured were preserved.

The process of monitoring the appearance and development of larvae, presence of pupae, the emergence of adults and deposition of egg rafts was repeated for the second, third and fourth generations when applicable. In each case, the date from the first appearance of egg rafts in any generation was used to estimate the duration of the life cycle time from egg-to-egg of each generation. Larvae were sampled and identified to species level to determine the population composition in each microcosm.

## Results

### Emergence timing

Of the 1228 emergent adults, all but four were identified as *C. obscuripes*. Counts of the numbers of emerged adult male and female *C. obscuripes* in each of the four microcosms are shown in Figure 1.

**Figure 1:**
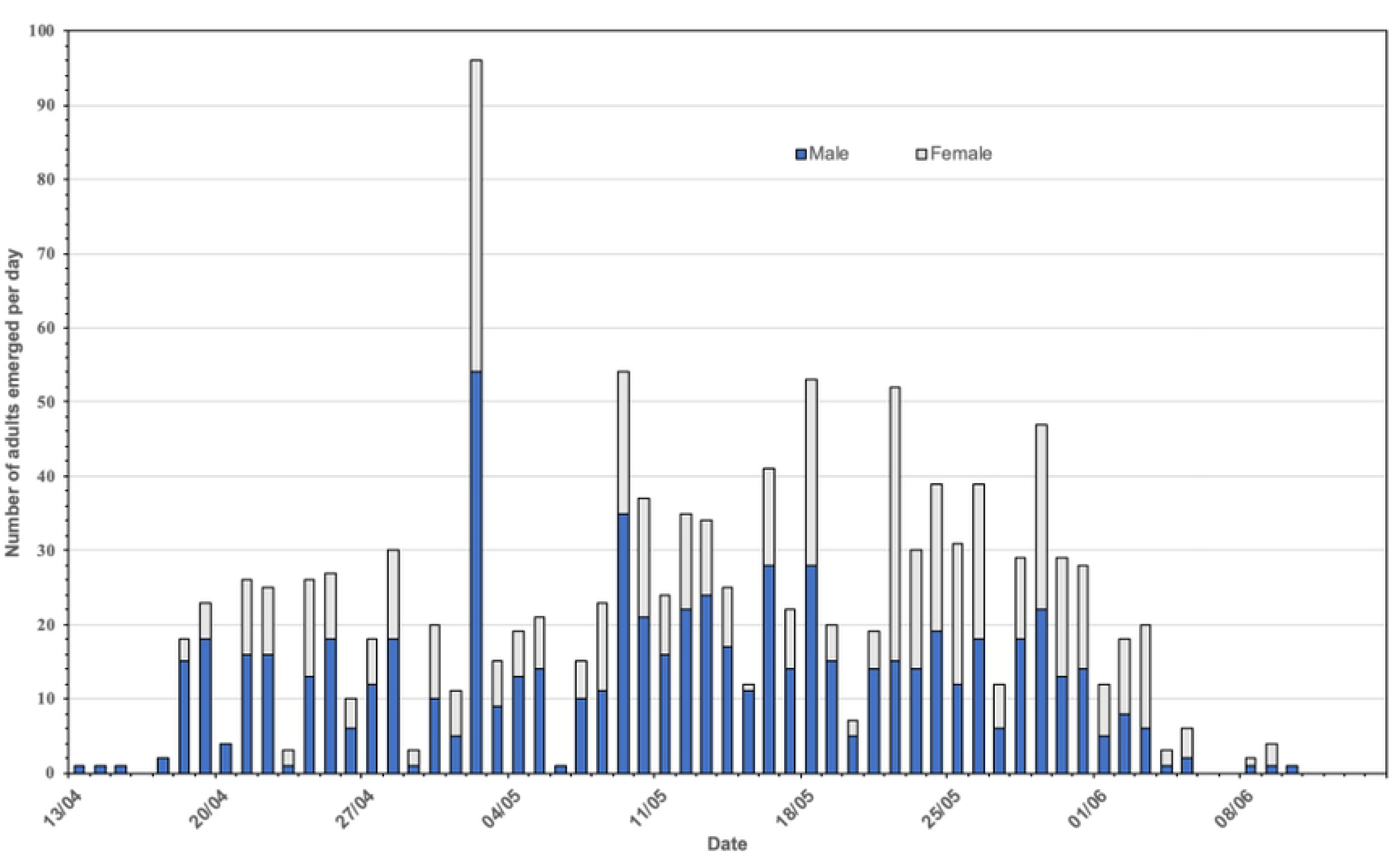
Emergence timing of male and female *Chaoborus obscuripes*.

Three specimens of *C. crystallinus* were found, the first (in Replicate 2) on 24 April 2017 (Day 25 post-initiation) and one each on 01 and 05 September 2017 (Days 154 and 158, respectively, both in Replicate 1). A single adult *Chaoborus* sp. sampled from Replicate 1 on 22 May 2017 (Day 53) could not be identified to species level. Therefore, of the successfully emerged adults, 99.68% were *C. obscuripes* and only 0.24% were *C. crystallinus*.

Males appeared to emerge slightly earlier than females (Figure 1). In the first week of emergence 42 males were recorded compared with 8 females. Before 22 May there were generally more freshly emerged males than females whereas the reverse was observed after this date.

Mean values for water temperature, dissolved oxygen, pH and conductivity measured weekly in each microcosm are summarised in Table 3. Water temperature was measured every 30 minutes throughout the study in one microcosm in replicate 1. The raw data for these environmental condition readings are available in a data file “Environmental Conditions for Emergence study” and have been summarised as daily maximum, minimum and mean values in Figure 2.

**Table 3:** Weekly water conditions during the *Chaoborus* emergence study

| **Date measured** | Mean Water Temp. (°C)<br>(Standard Deviation) | Mean Water pH<br>(Standard Deviation) | Mean DO<sub>2</sub> (%ASV*)<br>(Standard Deviation) | Mean Cond. (µS cm<sup>-1</sup>)<br>(Standard Deviation) |
| --- | --- | --- | --- | --- |
| 05-Apr | 13.50 (0.24) | 7.81 (0.04) | 92.98 (8.54) | 961.25 (9.18) |
| 12-Apr | 12.25 (0.13) | 8.17 (0.09) | 127.45 (18.54) | 854.00 (16.02) |
| 19-Apr | 9.20 (0.23) | 8.09 (0.11) | 124.23 (7.23) | 814.00 (5.72) |
| 27-Apr | 8.33 (0.10) | 8.71 (0.12) | 134.05 (2.82) | 766.00 (6.98) |
| 03-May | 11.00 (0.14) | 9.32 (0.07) | 138.875 (2.17) | 745.00 (4.55) |
| 11-May | 14.60 (0.61) | 9.74 (0.01) | 147.30 (4.20) | 733.25 (6.34) |
| 19-May | 15.35 (0.10) | 9.74 (0.05) | 133.95 (0.83) | 678.75 (8.18) |
| 26-May | 17.30 (0.32) | 9.78 (0.14) | 132.05 (7.27) | 683.25 (9.57) |
| 02-Jun | 19.83 (0.33) | 10.20 (0.35) | 152.78 (19.52) | 682.25 (10.69) |
| 09-Jun | 20.48 (0.26) | 10.38 (0.15) | 171.50 (6.88) | 637.25 (11.73) |
| 16-Jun | 21.68 (0.15) | 10.33 (0.02) | 178.58 (14.48) | 653.00 (11.92) |

\* ASV - Air Saturation Value

**Figure 2:**
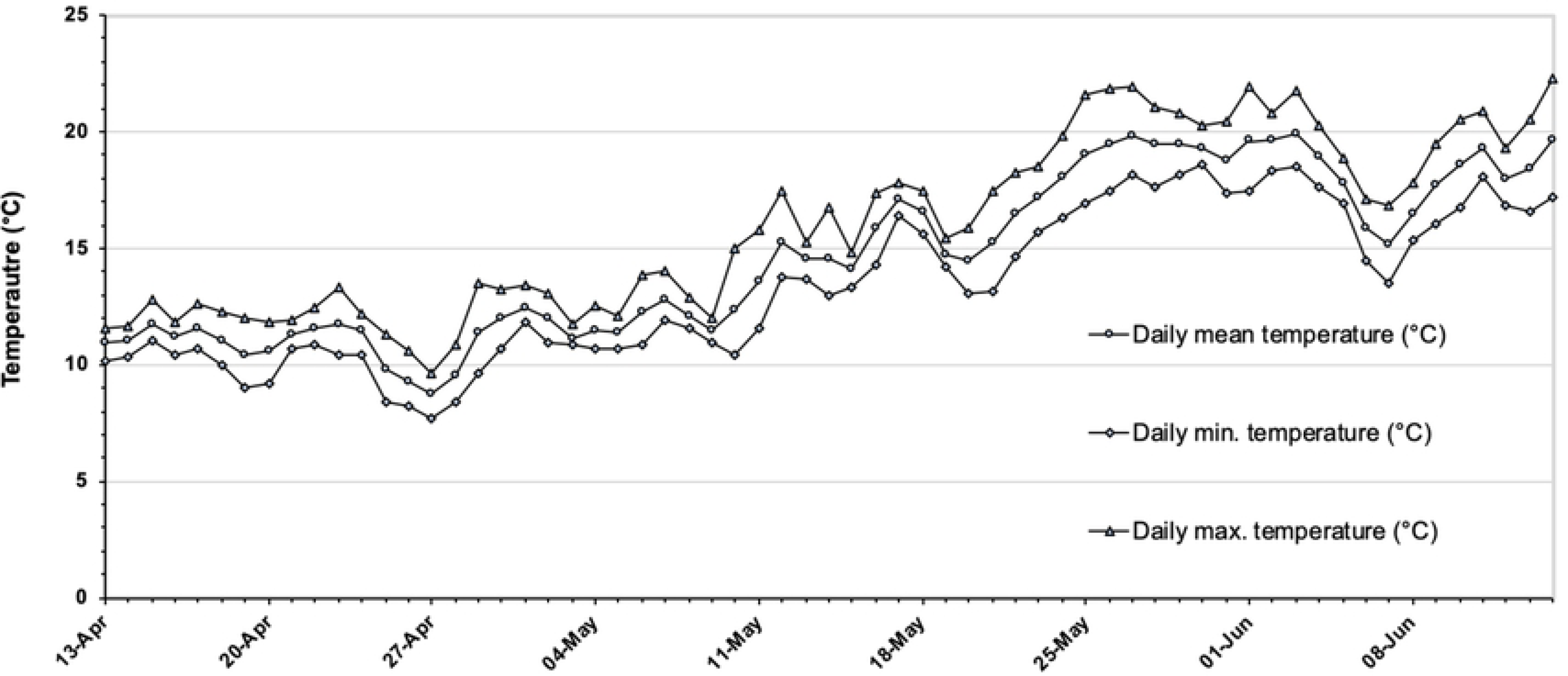
Daily maximum, minimum and mean water temperatures recorded in replicate 1 during the emergence study.

### Voltinism

Daily mean water temperatures, pH, dissolved oxygen and conductivity values recorded during the life cycle trial are presented in Table 5. The development of *Chaoborus* from the initial egg rafts introduced into each unit are presented for each replicate in Figure 3. Egg rafts added to the first microcosms of replicates 1, 2, 4 and 5 failed to establish at the first attempt and replicates 1, 2 and 5 were re-initiated at intervals as fresh egg rafts became available. The re-initiated replicates 1 and 5 both progressed through to a fourth generation. These two replicates were found to contain both *C. obscuripes* and *C. crystallinus* in their respective first units although the larvae sampled from units 2 – 4 in both replicates were all *C. crystallinus*. In both replicates, *C. obscuripes* were only found in July, while the specimens of *C. crystallinus* were only found in October. Replicate 4 could not be re-initiated with egg rafts as none were available at a suitable time. Monitoring of this unit showed that no larvae were present at any time, confirming the effectiveness of the systems in preventing immigration of *Chaoborus* spp. from outside. The re-initiation of replicate 2 produced only two generations which, when sampled in October, were found to consist only of *C. crystallinus*.

**Figure 3:**
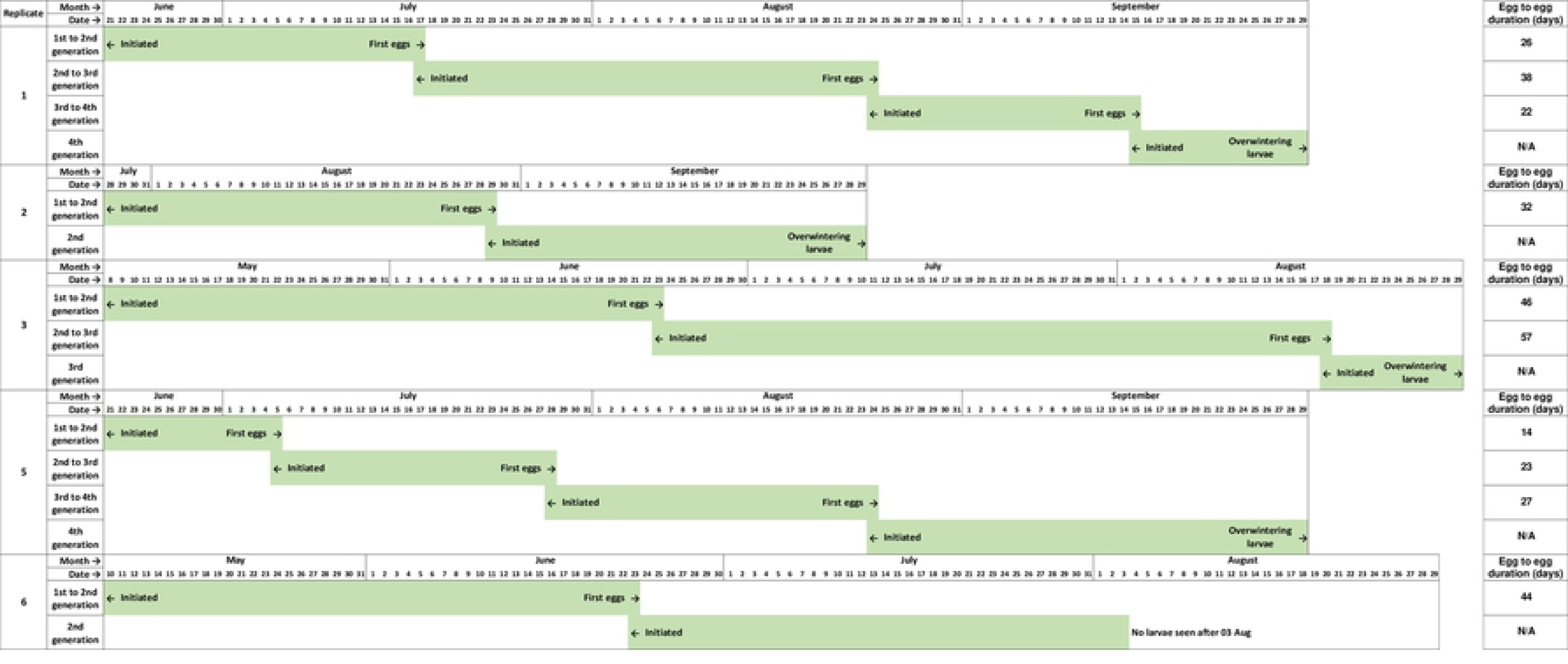
Development of *Chaoborus* from introduced egg rafts in each microcosm that sustained populations.

Populations of *Chaoborus* in replicates 3 and 5 progressed through to a third and fourth generations respectively. Larvae sampled from the first unit in July were found to be *C. obscuripes*, although when re-sampled in October, both *C. obscuripes* and *C. crystallinus* were found to be present. Generations 2 and 3, both sampled in October, were found to consist only of *C. crystallinus*. Replicate 6 also did not require re-initiation but as the second unit did not establish successfully, larvae were not sampled from Unit 1 until October, in order to give the maximum opportunity for more egg deposition to restart Unit 2. In practice, a second production of egg rafts did not occur in Unit 1 and therefore, Unit 2 could not be re-started. Only six late-instar larvae remained in Unit 1 by the October sampling and all were found to be *C. obscuripes*.

Minimum egg-to-egg times for *C. crystallinus* are summarised in Table 4 and ranged from 14 days (Replicate 5, Unit 1) to 56 days (Replicate 3, Unit 2). As only *C. crystallinus* was found in the second, third and fourth generations of any replicate, it is not possible to draw any conclusions regarding the egg-to-egg timings for *C. obscuripes*. The shortest generation time of 14 days occurred when the water temperature was at its highest (Fig. 3) in late June and early July. However, the longest observed development time of 56 days also spanned this period so other variables such as prey density are clearly involved in determining *Chaoborus* development rate.

**Table 4:** Egg to egg development times of *Chaoborus crystallinus*

|  | Unit 1 |  | Unit 2 |  | Unit 3 |  |
| --- | --- | --- | --- | --- | --- | --- |
|  | Dates / Duration (days) |  | Dates / Duration (days) |  | Dates / Duration (days) |  |
| Replicate 1 | 21 June-17 July | (26) | 17 July- 24 Aug. | (38) | 24 Aug-15 Sept* | (22) |
| Replicate 2 | 28 July - 29 Aug. | (32) | No successful second generation |  | No successful third generation |  |
| Replicate 3 | 8 May-23 June | (46) | 23 June - 18 Aug. | (56) | No successful third generation |  |
| Replicate 5 | 21 June- 5 July | (14) | 5-28 July | (23) | 28 July - 24 Aug.* | (27) |
| Replicate 6 | 10 May-23 June | (44) | No successful second generation |  | No successful third generation |  |
| Mean | 32.4 |  |  | 39.0 |  | 24.5 |
| Overall mean | 32.8 (S.D.13.0 days) |  |  |  |  |  |

Although environmental conditions were recorded in all replicates during the study, they did not differ significantly for any of the measured variables. The water temperature, dissolved oxygen levels, pH and conductivity are presented in Table 5. The presence of the mesh enclosures had negligible impact on water temperature in the microcosm. Temperatures in an open microcosm were compared to those in an enclosed one and found to be very similar throughout the study.

**Table 5:** Environmental conditions in microcosms in the life cycle study

| Date | Mean Temp. (°C) | STDEV of Temp | Mean pH | STDEV of pH | Mean DO<sub>2</sub> (%) | STDEV of DO<sub>2</sub> | Mean Cond. (µS cm<sup>-1</sup>) | STDEV of Cond | (n) |
| --- | --- | --- | --- | --- | --- | --- | --- | --- | --- |
| 25-Apr | 13.5 | n/a | 8.1 | n/a | 134.6 | n/a | 802 | n/a | 1 |
| 27-Apr | 9.2 | n/a | 8.2 | n/a | 114.9 | n/a | 785 | n/a | 1 |
| 03-May | 11.0 | 3.04 | 8.3 | 0.24 | 102.7 | 10.32 | 818 | 8.49 | 2 |
| 08-May | 13.4 | 0.14 | 8.4 | 0.20 | 103.6 | 7.64 | - | - | 2 |
| 10-May | 16.8 | 0.35 | 8.5 | 0.13 | 107.7 | 13.15 | - | - | 2 |
| 11-May | 15.5 | 0.43 | 8.4 | 0.18 | 107.3 | 11.65 | 793 | 44.08 | 6 |
| 19-May | 15.3 | 0.18 | 8.6 | 0.21 | 112.6 | 11.80 | 739 | 45.68 | 6 |
| 26-May | 17.6 | 0.48 | 8.8 | 0.56 | 122.1 | 18.77 | 724 | 51.04 | 6 |
| 02-Jun | 18.8 | 0.40 | 9.4 | 0.92 | 141.5 | 40.68 | 713 | 65.88 | 6 |
| 09-Jun | 19.3 | 0.60 | 10.0 | 0.87 | 162.3 | 23.52 | 670 | 54.65 | 6 |
| 16-Jun | 21.8 | 0.38 | 10.2 | 0.31 | 172.4 | 19.28 | 675 | 44.48 | 6 |
| 21-Jun | 20.4 | 0.19 | 10.0 | 0.24 | 156.4 | 18.23 | 676 | 40.69 | 6 |
| 23-Jun | 23.2 | 0.85 | 10.0 | 0.09 | 156.5 | 13.44 | - | - | 2 |
| 28-Jun | 17.7 | 0.12 | 10.1 | 0.34 | 156.1 | 19.33 | 661 | 40.05 | 8 |
| 07-Jul | 25.2 | 0.49 | 10.2 | 0.28 | 196.7 | 32.63 | 674 | 49.94 | 7 |
| 14-Jul | 19.2 | 0.27 | 10.3 | 0.22 | 170.4 | 18.59 | 688 | 109.07 | 10.00 |
| 21-Jul | 21.3 | 0.20 | 10.2 | 0.26 | 205.4 | 19.18 | 661 | 42.34 | 9.00 |
| 28-Jul | 19.0 | 0.30 | 10.1 | 0.39 | 160.5 | 26.67 | 627 | 42.30 | 12.00 |
| 02-Aug | 18.3 | 0.56 | 10.14 | 0.47 | 165.2 | 26.30 | 645 | 54.24 | 12.00 |
| 11-Aug | 19.5 | 1.02 | 10.12 | 0.44 | 156.9 | 20.84 | 601 | 40.65 | 12.00 |
| 18-Aug | 19.9 | 0.39 | 10.00 | 0.42 | 163.2 | 14.88 | 599 | 46.70 | 12.00 |
| 24-Aug | 19.4 | 0.07 | 9.8 | 0.39 | 130.8 | 45.33 | - | - | 2 |
| 25-Aug | 17.1 | 0.48 | 9.7 | 0.36 | 123.2 | 22.14 | 583 | 49.29 | 14.00 |
| 29-Aug | 22.6 | n/a | 9.2 | n/a | 139.6 | n/a | - | - | 1 |
| 06-Sep | 16.8 | 0.21 | 9.7 | 0.39 | 136.9 | 20.62 | 588 | 58.84 | 15.00 |
| 13-Sep | 13.9 | 0.24 | 9.4 | 0.55 | 111.2 | 15.97 | 588 | 58.55 | 15.00 |
| 15-Sep | 14.6 | n/a | 7.5 | n/a | 96.8 | n/a | - | - | 1 |
| 20-Sep | 12.3 | 0.13 | 9.6 | 0.54 | 113.3 | 12.43 | 577 | 57.73 | 15.00 |

## Discussion

Emergence of adult *Chaoborus* spp. from overwintered larvae in South Eastern England commenced on 13 April 2017 (14 days after being introduced to artificial microcosms, Day 0) and peaked on 2 May 2017 (Day 33). The majority of emergence was completed by early June (the last occasion when emergence of *Chaoborus* spp. occurred in all four replicates was 3 June 2017, Day 65). Of the 2027 fourth instar larvae introduced into the microcosms, the mean emergence success was 60.9%, ranging from 51.4% in Replicate 4 to 66.2% in Replicate 1. As indicated in (22), it is likely that carnivory was responsible for the emergence success being less than 100%. All except four of the emergent adults were *C. obscuripes*, which was consistent with the populations of *Chaoborus* spp. known to inhabit the source pond in previous years. These results show that the emergence of adult *C. obscuripes* originating from the pupation of post-overwintering fourth instar larvae took place over a clearly defined period between mid-April and early June. Males appeared to emerge slightly earlier than females (Figure 1). In the first week of emergence 42 males were recorded compared with 8 females.

The results of the voltinism study showed that *C. crystallinus* produced up to four discrete generations within the experimental period. Since two replicate microcosm groups exhibiting four generations were both re-initiated several weeks after the season’s egg deposition commenced, one more generation may have been possible. There may be some differences in developmental times or reproductive success between species since from the second generation onwards populations were dominated by *C. crystallinus*, despite the presence of *C. obscuripes* in the first generations. It is possible that the test units or their conditions may have been favourable for oviposition by *C. crystallinus* but not by *C. obscuripes*. In "the wild" C. *obscuripes* is almost always associated with larger bodies of water (large ponds, small lakes and reservoirs).

The time for development from egg-to-egg ranged from 14 days to 56 days with a mean of 32.8 days (Standard Deviation 13.0 days) indicating a high degree of phenotypic plasticity in *C. crystallinus* life history strategies. Given that environmental conditions within the microcosms were all very similar it seems most likely that differing levels of prey density contributed to the wide range of development times. Prey abundance and diversity was not determined as the influence of prey availability was outside the scope of this study. However, this warrants further investigation as feeding is clearly a significantly influential factor in development times.

This study confirmed that under temperate conditions *C. crystallinus* exhibits a multi-voltine life history.

